# Genome mining of *Streptomyces* for the discovery of low-resistance antibiotics

**DOI:** 10.1101/2023.12.28.573565

**Authors:** Rosete Ambriz Sergio Antony, Alejo Hernández Moisés Alejandro, Sánchez-Cruz Norberto, Ceapă Corina-Diana

## Abstract

2.

Antimicrobial resistance is considered one of the top ten global health crises, which requires discovering and developing antibiotics with low resistance. Historically, Streptomyces bacteria are well-known for adapting to their complex environment by accumulating numerous clusters of specialized metabolites, some of which remain inactive in laboratory settings. The genomic revolution significantly increased their potential, complementing laboratory practices, and allowing for the discovery of antimicrobials associated with this biosynthetic machinery that are not constitutively produced. In the current study, we used the latest bioinformatics analyses to improve upon these predictions for Streptomyces genomes, and to identify low-resistance novel antimicrobial candidates. Our integrated pipeline used antiSMASH, BiG-SCAPE/EFI-EST, BiG-FAM, and additional tools and identified 326 novel BGCs and 67 peptides with antimicrobial potential. Further exploration of ribosomally synthesized and post-translationally modified peptides (RiPPs) revealed diverse chemical structures and suggested new mechanisms of action. Artificial intelligence platforms, such as MACREL, predicted the antimicrobial activity of the identified peptides, offering a comprehensive strategy for discovering bioactive compounds. Their low-resistance potential was estimated on a case-by-case basis. This study showcases the extensive genomic potential of Streptomyces, providing valuable insights for future antibiotic discovery efforts.

**Graphical abstract:** 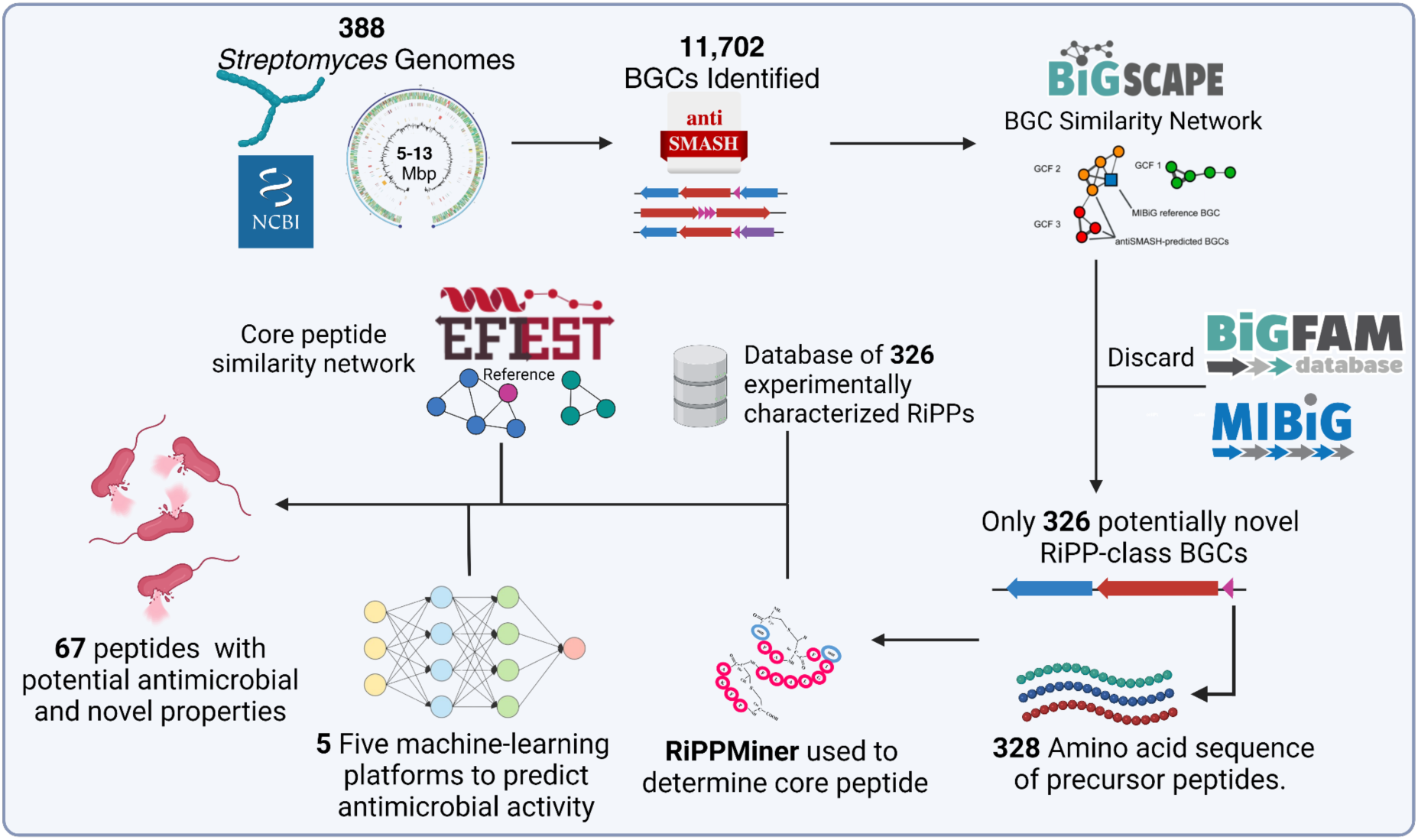

**Impact statement:** Exploring the potential applications of microorganisms is essential in fields like medicine, ecology, and biotechnology. Genomics is an invaluable tool that can uncover secondary metabolites with various applications in these fields. To this end, we conducted a detailed analysis of 388 complete genomes of Streptomyces, with a focus on identifying new biosynthetic gene clusters (BGCs) and antimicrobial peptides. Our analysis involved the use of various tools, including AntiSMASH, BiG-SCAPE, and reference databases like MiBIG and BiG-FAM, to unravel the diversity of these biosynthetic systems. We established a manually curated database and a robust pipeline for identifying valuable compounds, which allowed us to prioritize 326 BGCs with novel and diverse biosynthetic machinery. These unique insights into the genomic richness of Streptomyces will serve as a valuable guide for future antibiotic discovery efforts, facilitating the selection of strains that can produce new natural products. Moreover, our approach also helped us identify 67 potential antimicrobial peptides, including some belonging to the newly discovered Class V lanthipeptides, highlighting the diversity and promise of these compounds. Our work offers a glimpse into the potential of genomics in identifying important molecules and provides a framework for future studies in this field.

**Data summary:** Big tables and special files can be found in Zenodo, under the following link In there, the following objects are included:

- Supplementary Table 1: a spreadsheet with nine tables: (Table S1) Accession numbers and genomic characteristics of the genomes used in this study; (Table S2) summary of predicted Biosynthetic Gene Clusters (BGCs) for each genome; (Table S3) summary of the distribution of BGC types present in *Streptomyces* genomes; (Table S4) IDs and strains of MiBIG BGCs; (Table S5) All information regarding the database of peptides with antimicrobial activity; (Table S6) Data of BGCs recovered after the analysis of MiBIG and BiG-FAM; (Table S7) Complete and potentially novel BGC data used for the creation of the SSN with EFI-EST; (Table S8) Data predicting potential antimicrobial activity using MACREL v1.2.0; (Table S9) Data of predicted peptides with positive antimicrobial activity. These data were used to create the graph showing the distribution of RiPPs classes within these peptides.
- Supplementary Data 1: a folder containing all .gbk files for the BGCs predicted by antiSMASH and MiBIG. These files were used as input for BiG-SCAPE (https://drive.google.com/file/d/1VGWJo9rsF429IUFf2g-tqs_NfBicNX91/view?usp=drive_link).
- Supplementary Data 2: amino acid sequences of the mined and reference peptides (from the database built into this work) in .fasta format.
- Supplementary Table 2: a spreadsheet with five tables: Data from the evaluation of the prediction of antimicrobial activity with five different AIs through five comparisons of the created database (Table S1-Macrel); (Table S2-CAMPR3); (Table S3-AMP Scanner); (Table S4-AmPEP); (Table S5-AL4AMP).
- Supplementary Figure 1: results of the predictive performance analysis (script: https://github.com/Bert-IQ/ripps-classification).

## 4. Introduction

Given the current spread of antimicrobial resistance, considered one of the top 10 global public health threats by the WHO [1], there has been an increase in severe infections caused by antibiotic-resistant bacteria. These bacteria do not respond to available treatments, primarily due to the misuse and overuse of antimicrobials [2]. Consequently, discovering new antibiotics, particularly those with novel modes of action, is essential to preventing future catastrophic pandemics [3].

One of the strategies for antibiotic discovery is based on natural products, as they have been the source of most clinically used antibiotic classes [4]. Actinobacteria, a group of filamentous Gram-positive bacteria with linear genomes and a high guanine and cytosine content (>70%) are the main source of natural antibiotics [5]. Among them, *Streptomyces*, the largest genus of actinobacteria, is known for producing many specialized metabolites with applications in agriculture, biotechnology, and medicine [6]. Despite not being essential for growth, these secondary metabolites (SMs) play crucial roles in the organism’s survival, exhibiting antiviral, antifungal, anticancer, immunosuppressive, and antibiotic properties [7,8].

However, *Streptomyces’s* genetic potential has been understudied because many SMs are “cryptic”, meaning they are encoded in the bacterial genome but not expressed under laboratory conditions [7,9]. The synthesis of secondary metabolites (SMs) in bacteria is a complex process that is regulated by intricate networks under the influence of biotic and abiotic factors in the organism’s natural environment. Furthermore, the synthesis of SMs occurs in conjunction with multiple secondary metabolic pathways, which can limit yields and complicate identification. SMs are produced by complex multienzyme systems encoded by secondary metabolite biosynthetic gene clusters (smBGCs). Based on the central gene, smBGCs are classified into five categories: non-ribosomal peptide synthetases (NRPS), polyketide synthases (PKS), hybrids of non-ribosomal peptide synthetases and polyketide synthases (NRPS-PKS), terpenes, and post-translationally modified peptides (RiPP). However, the expression of these smBGCs groups is challenging, and only a small fraction of secondary metabolites has been produced under laboratory conditions, with over 90% of smBGCs remaining unexpressed and characterized [10-12]. It is imperative to also consider that most investigations in this domain rely on homology. As a result, any gene that represents a particular biosynthetic class that is not incorporated in the databases will not be detected. Nonetheless, recent advancements in both analysis pipelines and databases have propelled this field forward, leading to the detection of novel chemical entities and bioactivities. These developments have played a crucial role in expanding our understanding of the molecular mechanisms underlying biological processes and in the identification of new targets for drug discovery.

In this way, the biosynthetic potential of these microorganisms has been underestimated. However, with the rapid advancements in next-generation sequencing (NGS) technology, there has been a high availability of freely accessible genomic data [10]. This has shifted the traditional approach to antimicrobial discovery, which relied on chemical identification through phenotypic detection and often led to rediscovery [13]. Instead, genome mining has emerged as a reverse approach, allowing the determination of biosynthetic gene clusters (BGCs) for known secondary metabolites from genomes published in databases. Additionally, the chemical structures of their products can be predicted to some extent based on the analysis and biosynthetic logic of enzymes encoded in a BGC and their similarity to known homologs [14]. This approach has unveiled the metabolic capabilities of model organisms producing natural products, such as *Streptomyces coelicolor*, revealing the biosynthetic potential within this genus [15].

Genome mining has been developed thanks to computational tools for BGC identification, such as antiSMASH [16,17] and PRISM [18], and the prediction of the chemical structures of their products. Furthermore, comparative analyses of predicted BGCs with biochemical reference data from databases like Minimum Information about a Biosynthetic Gene cluster (MIBiG) have strengthened this *in silico* approach [19].

The availability of vast genetic information has enabled systematic investigations into the biosynthetic potential of large taxonomic groups of organisms [20-23]. These comprehensive analyses allow the straightforward identification of thousands of BGCs with varying levels of similarity, spanning from widely distributed homologs related to the production of known molecules to rare or unique gene clusters encoding enzymes and synthesis pathways that are still unknown [24]. To map and prioritize this complex biosynthetic diversity, tools have been developed that compare architectural relationships between BGCs in sequence similarity networks and group them into Gene Cluster Families (GCFs). BGCs within the same GCF are linked to very similar types of natural products [25]. However, these models exhibit low confidence in accurately measuring the similarity between complete and fragmented gene clusters, a phenomenon common in metagenomic projects. Additionally, the lack of integration among these tools into a single comprehensive methodology has hindered widespread adoption by the scientific community, impeding large-scale advancements in the discovery of novel bioactive natural products.

This study provides an optimized computational workflow that integrates various bioinformatics tools to prioritize the discovery of potentially novel antimicrobial secondary metabolites from complete sequences of the *Streptomyces* genus.

## 5. Methods

### Genome Selection

Complete *Streptomyces* genomes (as of September 2022) were downloaded from the National Center for Biotechnology Information (NCBI) in FASTA format. After manual curation, removing plasmid sequences, and repeated sequences, 388 complete genomes were included. Accession numbers and genomic information (genome size, number of BGCs, chromosome type) are shown in Supplementary Table 1. All graphics were created with Graph Pad Prism 8.

### BGC Prediction

Biosynthetic gene clusters were predicted using antiSMASH (Antibiotics and Secondary Metabolite Analysis Shell, v6.1.1), which identifies BGCs using a Hidden Markov Model based on multiple alignments of experimentally characterized protein sequences or protein domains [16]. The detection setting was “relaxed,” and all permitted options for analysis were enabled. The 388 *Streptomyces* genomes were used in FASTA format. By enabling the MIBiG cluster comparison option, antiSMASH compared the predicted BGCs with MIBiG’s BGCs, selecting BGCs with a similarity degree of 75% or more to the predicted BGCs to build a BGC Similarity Network. Descriptors (location, type, and comparison with MIBiG) of predicted regions for each sequence are shown in Supplementary Table 1.

### BGC Similarity Network

The Genbank files of 11,703 BCG derived from antiSMASH analysis and 203 BGC of reference from MIBiG database were used as input to build a BGC similarity network using BiG-SCAPE 1.1.2 [24] and in that way thereby cluster BGCs into CGF. Two different cutoff values, 0.3 and 0.6 in raw_index was tested to define network construction. However, as the sequences of many BGCs are very similar, we used the cutoff value of 0.3 for the final network (supplementary Data 2). The comparison was performed jointly for the major BGC classes using the -hybrids-on parameter. The obtained similarity networks were visualized using Cytoscape 3.9.1 [28].

### Prioritization of Potentially Novel BGCs

#### Comparison with Reference Databases (MiBIG and BiG-FAM)

Starting from the similarity networks obtained with a cutoff value of 0.3, CGFs that exhibited a MIBiG BGC were discarded. One representative BGC was chosen for each CGF and inputted into the BiG-FAM database, by using the ID from each antiSMASH analysis. BiG FAM compares query BGCs with 1,225,071 known BGCs (grouped into 29,955 GCFs) based on similar protein domain architectures [29]. This allows for the degree of similarity of query BGCs with currently known bacterial BGCs (functionally pre-characterized) to be established. At the end of the analysis, CGFs were discarded when the representative showed a significant degree of similarity with CGFs reported on the platform. This study discarded query CGFs with values below 900 in the pairing distance with BiG-FAM CGFs [30].

#### Prioritization of Complete RiPPs BGCs

Manual analysis was conducted to assess the competence of the mined BGCs, inspecting the output of antiSMASH analysis for the presence of minimal biosynthetic machinery. For the RiPP class BGCs, central biosynthetic genes and the precursor peptide gene were considered minimal.

#### Analysis and Prioritization of Novel RiPPs

Subsequently, amino acid sequences of precursor peptides from RiPPs BGCs were mined and, together with experimentally characterized RiPPs from NCBI and RiPPMiner, were used to build a similarity network using the Enzyme Function Initiative-Enzyme Similarity (EFI-EST) web platform [31]. The following parameters were used: fasta, fasta header reading: enabled, protein family addition options: none, E-value: 5 (default), fragments: disabled. After the initial analysis, the alignment score threshold was set to 6, the sequence length restriction with default values, and neighborhood connectivity was disabled. The files were downloaded upon completion, and the obtained networks were visualized and modified using Cytoscape 3.9.1. In addition, a database of characterized RiPPs and their biological activity was manually built and curated (supplementary Table 1).

#### Antimicrobial activity pipeline development and validation

The determination of antimicrobial activity of peptides is a challenging and time-consuming task. Fortunately, there are numerous databases of peptides that provide experimentally determined antimicrobial activity [32]. This information has led to the development of machine learning tools that can predict the antimicrobial potential of new peptides. These tools are valuable in prioritizing the isolation and characterization of peptides that require experimental validation. While these tools have shown promising results in various benchmarking tests, their ability to predict the antimicrobial potential of Ribosomally synthesized and post-translationally modified peptides (RiPPs) has yet to be examined.

This study evaluates five different machine learning models, including MACREL [33], CAMPR3 [34], Antimicrobial Peptide Scanner vr.2 [35], AMPEP [36], and AL4AMP [37], for their ability to predict the antimicrobial potential of a database of RiPPs that have experimentally determined antimicrobial activity. The evaluation of these models is critical to determine their validity and reliability in predicting the antimicrobial activity of RiPPs. This study aims to provide insights into the applicability of machine learning models in predicting the antimicrobial potential of RiPPs and their potential in accelerating the discovery of novel antimicrobial agents.

The selection of machine learning models used in this study was based on a set of rigorous criteria. Firstly, only public, and freely accessible models were utilized. Secondly, the models were required to have accessible training databases. Thirdly, the prediction outputs of these models were expected to provide a probability score of the antimicrobial potential of the compounds, ranging from 0 to 1, rather than a simple binary label of “yes” or “no”. Finally, the models had to be available either as an online platform or as a standalone application, to enable their use in the study.

For all the Ribosomally synthesized and post-translationally modified peptides (RiPPs) in our assembled database, predictions were obtained using all the different models. However, only the predictions for RiPPs that were not included in the training dataset of each model were further analyzed. The performance of each model in correctly classifying the RiPPs was evaluated based on two metrics: the Area Under its Receiver Operating Characteristic (ROC) Curve and the area under its Precision-Recall Curve.

Given that these models were not explicitly trained to predict antimicrobial activity on RiPPs, their probability scores were not well-calibrated for such a task. Therefore, the precision and recall for the two best-performing models were examined at different threshold values of the probability score. This enabled the identification of an optimal cutoff for the classification of RiPPs, based on the probability scores.

The final candidate gene clusters were manually checked for transporters, other resistance genes, and horizontal gene transfer evidence (presence of insertion elements, transposons, and plasmids). When resistance genes were present, each gene was subjected to BLAST across the complete Prokaryote taxonomy. The distribution of resistance mechanisms is considered optimal when limited to specific narrow taxonomic groups and when no evidence of horizontal transfer can be detected.

## 6. Results and Discussion

### General Features of *Streptomyces* Genomes

In the present study, 388 complete genomes of the genus *Streptomyces sp*. from the NCBI database were included. The genome size ranges from 5.28 Mb for *Streptomyces* sp. HSG2 to 12.47 Mb for *Streptomyces rapamycinicus* NRRL 5491, with an average of 8.3 Mb. Despite the increased genomic information available for this genus, the genome size values are very close to those reported in previous metagenomic analyses [21,38,39].

Most of the analyzed genomes were linear, although circular genomes were identified, including *Streptomyces cyaneochromogenes* strain MK-45, *Streptomyces sp*. AMCC400023, *Streptomyces ferrugineus* strain CCTCC AA2014009, *Streptomyces sp*. NEAU-sy36, *Streptomyces lavendulae* subsp. *lavendulae* strain Del-LP, and *Streptomyces tendae* strain 139.

### Distribution and Diversity of Biosynthetic Gene Clusters (BGCs) in *Streptomyces*

The present study involved the analysis of predicted genomic regions of interest using antiSMASH v6.1.1 in relaxed mode, resulting in the identification of 11,702 colinear genetic regions (or 16,361 distinct BGCs) from 388 Streptomyces sp. genomes. Notably, multiple BGCs could be present within each region, but their separation was not considered due to the colocalization of some of the BGCs. This approach proved particularly relevant as colocalization can indicate the presence of hybrid clusters, such as the NRPS/T1PKS modular system, which is a common feature in bacterial systems [40]. Indeed, the most frequent hybrid BGC identified in this study was the NRPS/T1PKS module, as shown in Figure 1d. This finding confirms the high prevalence of such systems, which has been previously reported [41]. Moreover, identifying numerous other systems besides this module further underscores this observation.

**Fig.1.**
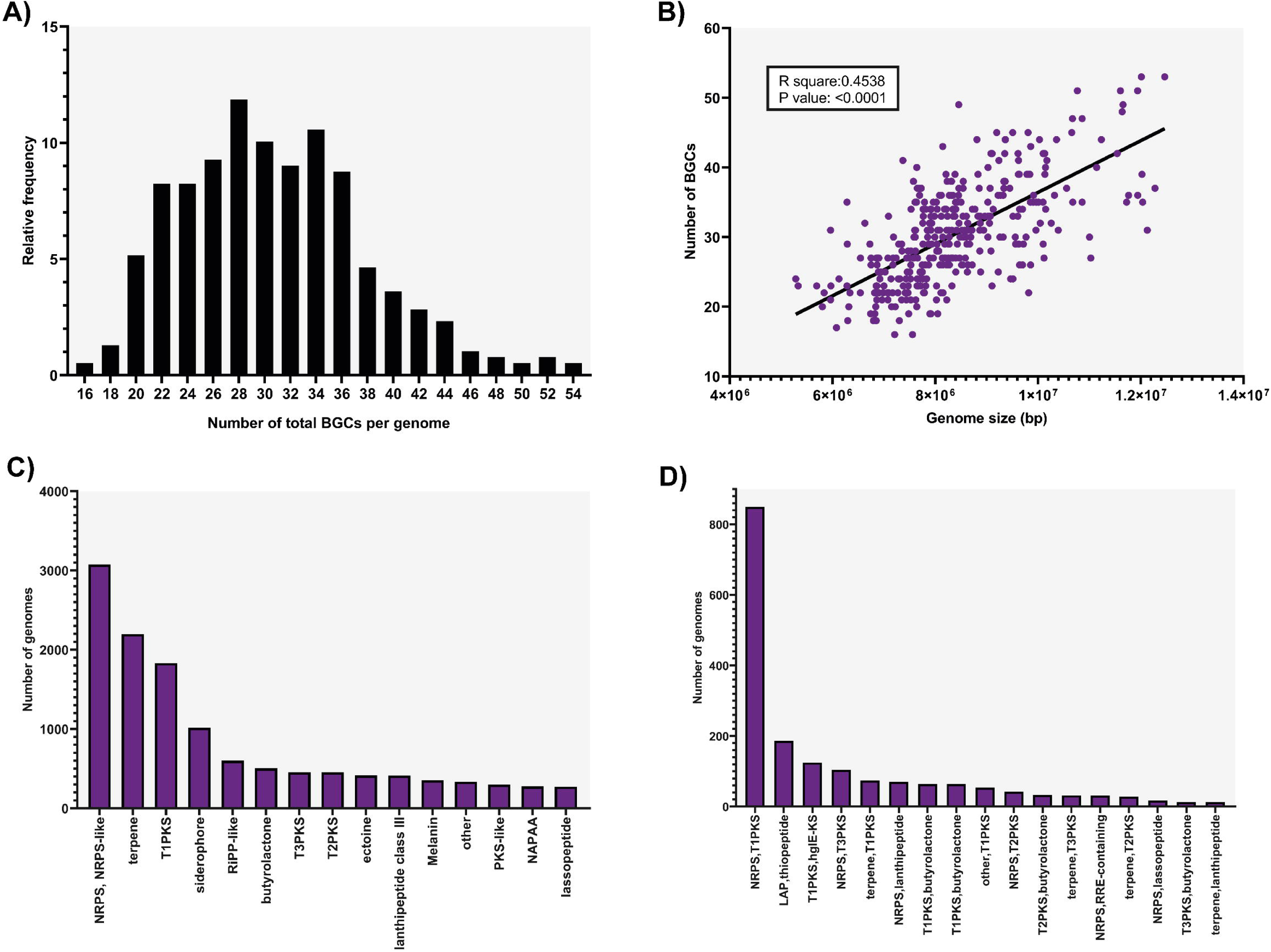
Frequency and diversity of BGCs. (a) Relative frequency distribution showing the total number of BGCs per genome. (b) Relationship between genome size and the total number of BGCs per genome. (c) Bar graph indicating the distribution of BGC classes in *Streptomyces* genomes by BGC type. (d) Bar graph indicating the major hybrid classes of BGCs.

Each genome presents 16 to 53 BGCs, consistent with the most recent metagenomic studies [38]. The strains situated at the extremities of the range are *Streptomyces sp*. WAC06273 strain WAC6273, with 16 BGCs, and *Streptomyces rapamycinicus* NRRL 5491, with 53 BGCs.

Fig. 1a shows a bell-shaped distribution relating the number of BGCs present in each genome by frequency, with 28 BGCs being the most frequent. This large number of identified BGCs per genome confirms the substantial potential of this genus. Additionally, a weak but significant positive correlation is observed between genome size and the number of BGCs per genome (r2 = 0.4538, p-value =<0.0001) (Fig. 1b), indicating an increase in the number of BGCs with a larger genome size, consistent with previous reports [41,42].

The considerable diversity of classes in *Streptomyces* genomes can be observed in Fig. 1c, representing 52 of the 70 classes of BGCs predicted by antiSMASH v6.1.1. For this analysis,

BGCs grouped in a single region were considered independent, resulting in 16,361 BGCs. Within these results, the NRPS class stands out with the highest distribution (18.79%), followed by terpenes (13.44%), T1PKS (11.19%), siderophores (6.20%), and RiPP-like (5.34%). These five classes are the most abundant in various analyses for the *Streptomyces* genus [24,41]. Additionally, the most common hybrid BGCs were NRPS/T1PKS (850 CGB), LAP/thiopeptide (186 BGC), and T1PKS/hglE-KS (125 BGC) (Fig. 1d). The complete numbers and types of BGCs predicted by antiSMASH for each genome are available in Supplementary Table 1.

To obtain a more detailed description and evaluation of the diversity present in Streptomyces Biosynthetic Gene Clusters (BGCs), a Sequence Similarity Network (SSN) was constructed using the BiG-SCAPE platform. This network clusters BGCs with similar biosynthetic machinery into Gene Cluster Families (GCF), enabling a biosynthetic functional classification. In cases where BGCs do not form a GCF, it suggests a potential lack of “function,” whether due to incorrect BGC prediction, incomplete BGCs, or an uncharacterized function due to significant divergence from known ones. Nodes that were not connected were therefore excluded from the GCF definition.

The SSN was based on similarities between BGCs for the RiPPs class, which showed 2519 BGCs. These were grouped into 381 networks (clans) containing more than two BGCs and 362 singleton BGCs. Most GCFs were formed by a single subclass of RiPPs, as shown in Figure 2. This SSN demonstrates a high diversity in the available biosynthetic machinery in Streptomyces, even when BGCs are grouped. Through this approach, the functional classification of BGCs can be better understood, which is essential for discovering novel natural products with potential pharmaceutical applications.

**Fig.2.**
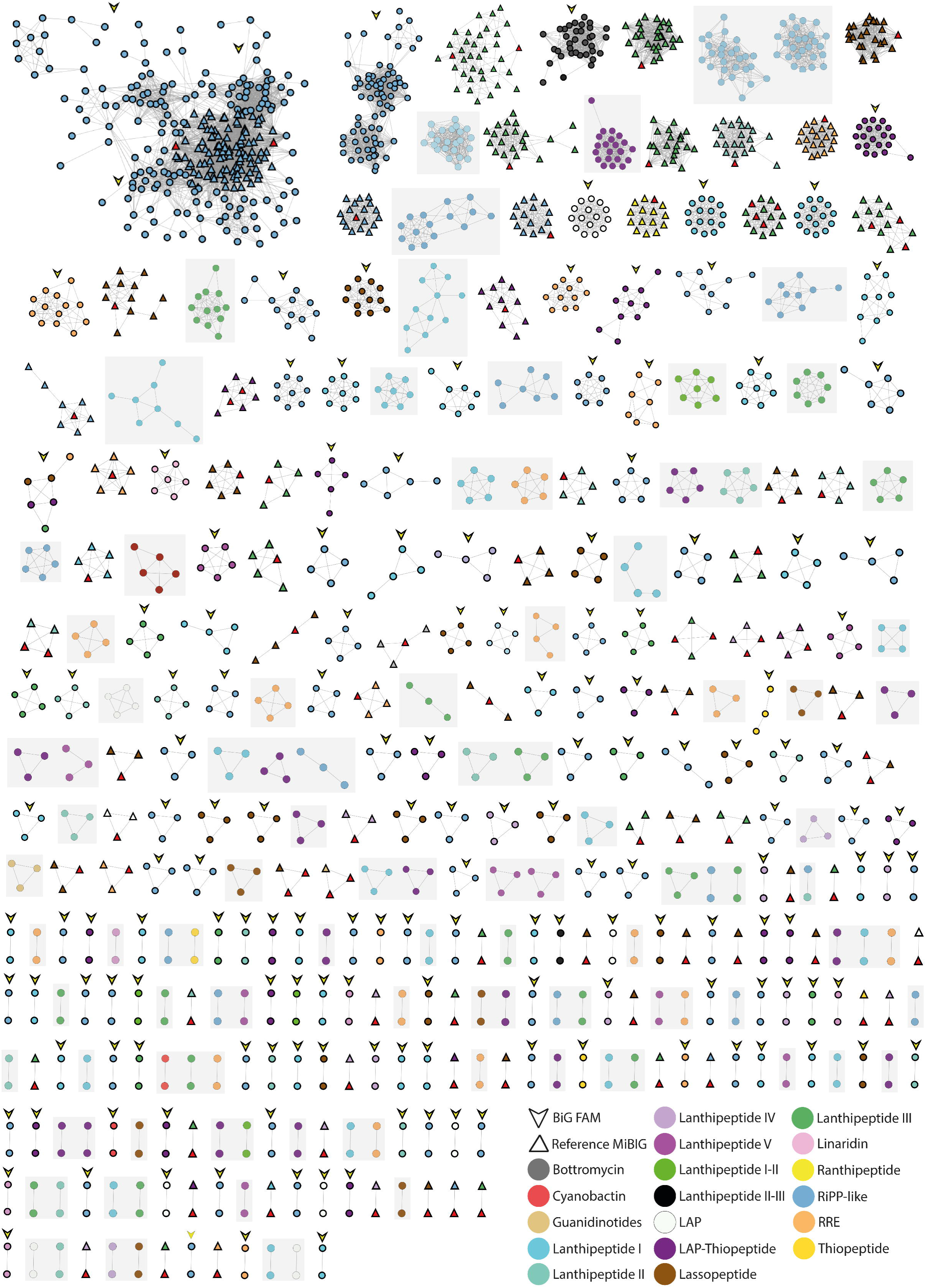
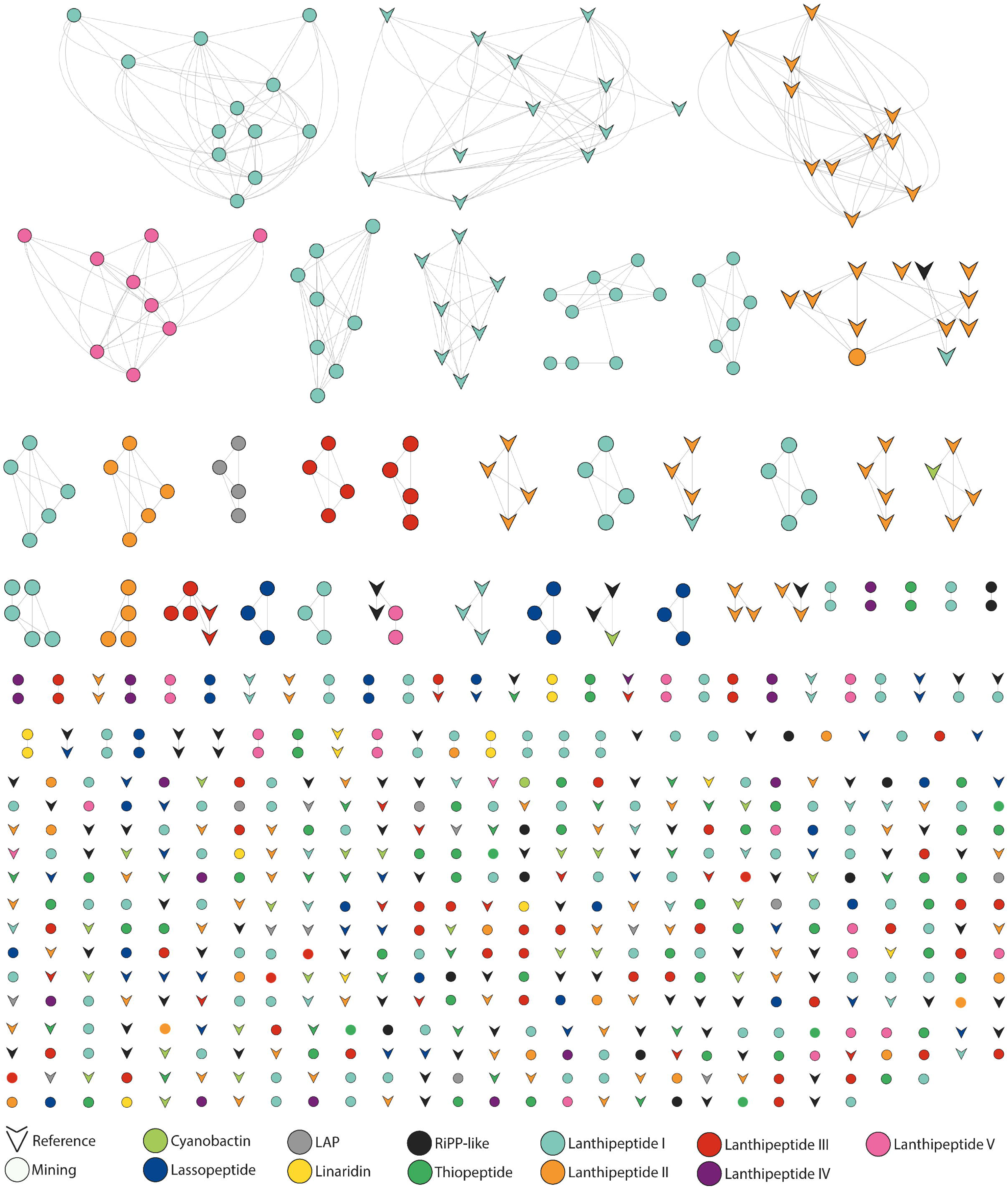
Similarity network of the RiPP class. The analysis was performed using as input the 11702 regions obtained from the antiSMASH v6.1.1 analysis and the BiG-SCAPE with a cutoff value of 0.3.

### Prioritization of Potentially Novel RiPP BGCs

One of the major challenges faced in drug discovery is the effective prioritization of strains that exhibit a higher potential for producing novel chemical compounds. A proposed methodology to address this issue involves using two reference databases - MiBIG and BiG-FAM. The MiBIG database assesses novelty at the biosynthetic gene cluster (BGC) level, while the BiG-FAM platform operates at the chemical-genetic functional annotation (CGF) level. Connecting these databases with the Secondary Metabolite Biosynthetic Gene Clusters Analysis Pipeline (BiG-SCAPE) makes it possible to perform large-scale analyses, which is facilitated by the high availability of genomic information through sequencing platforms. Such analyses are typically infeasible when using these tools in isolation.

### Comparison of SSN with Reference Databases (MiBIG and BiG-FAM)

Using two novelty criteria, 484 BGCs with potentially novel genetic machinery were identified. The first criterion involved the construction of SNN using the predicted BGCs by antiSMASH and 204 reference MiBIG BGCs, which exhibited at least 75% similarity with some of the BGCs predicted. This approach led to the exclusion of 75 CGFs that contained at least one MiBIG’s BGC, thereby obtaining BGCs with potential novelty.

The second criterion was to evaluate representative BGCs for each CGF without MiBIG BGC within them using the BiG-FAM platform. It allowed the comparison with 29,955 GCFs. This facilitated the determination of the degree of similarity between the query BGCs and currently known bacterial BGCs. The evaluation of potentially novel GCFs exceeding a pairing distance greater than 900 resulted in identifying 99 GCFs with the potential to encode novel natural products. The rigorous methodology described enabled the prioritization of novel BGCs, excluding approximately 70% of the predicted CGFs as being identical or closely related to previously characterized products, at the same time, validating the methodology.

### Prioritization of Complete RiPP BGCs

The viability of candidate Biosynthetic Gene Clusters (BGCs) is unknown as their biosynthetic competence has not yet been evaluated. Therefore, it is imperative to identify and select BGCs with minimal biosynthetic machinery. In the case of RiPPs, the minimum machinery required involves the presence of the precursor peptide gene and central modifying enzymes. It has been discovered that many natural product BGCs do not harbor a complete set of necessary genes to produce specialized metabolites and must use enzymes encoded elsewhere in the genome for biosynthesis [43]. For RiPPs, the protease gene is frequently absent within the BGC or included as a module within the transporter gene, making the prediction of this class of BGCs challenging. Therefore, this gene is not considered within the minimal machinery required to identify complete BGCs [44].

Following an analysis of the outputs from antiSMASH for each BGC, 326 BGCs with novel biosynthetic machinery for producing specialized metabolites were identified. Notably, the primary reason for discarding incomplete BGCs was the absence of the gene responsible for encoding the precursor peptide, which is a crucial criterion in predicting RiPPs, particularly for new classes [45,46].

### Analysis and Prioritization of Novel Peptides

The primary aim of the methodology described here was to prioritize novel Biosynthetic Gene Clusters (BGCs), with a specific focus on Ribosomally synthesized and post-translationally modified peptides (RiPPs). In this class of BGCs, the precursor peptide-encoding genes directly impact the structure of the produced metabolite. However, due to their small size, these genes have low significance in the sequence comparisons used at the BGC level, often resulting in the overlooking of significant similarities with reference genes. Therefore, additional analysis is necessary to prioritize potentially novel specialized metabolites at the single-gene level. The EFI-EST platform was utilized to generate a Self-Organizing Neural Network (SNN) of single amino acid sequences to address this issue.

To construct the SNN, predicted sequences and reference sequences were input into the EFI-EST platform. A similarity value of 35% was chosen, and the resulting network was visualized with Cytoscape. As illustrated in Figure 3, the resulting network indicates that most central peptides are not grouped into any family, suggesting a vast chemical diversity of specialized metabolites in the Streptomyces genus. Moreover, a limited number of peptides were grouped with experimentally characterized peptides, indicating that most peptides are potentially novel. In total, 328 potentially novel peptides with minimal biosynthetic machinery for production were obtained through this method.

**Fig.3.**
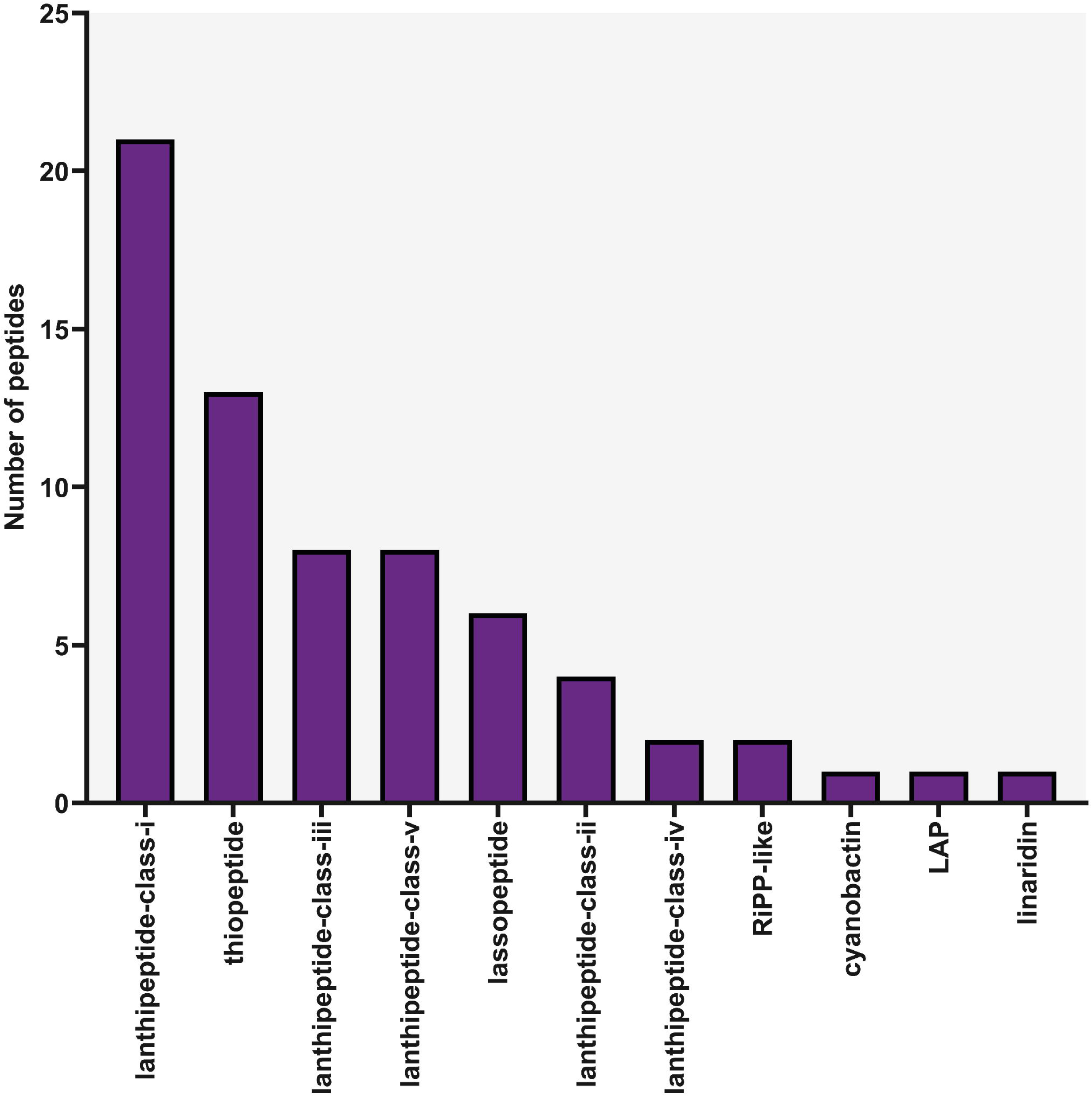
Prioritization of novel peptides of the RiPP class. SSN of the core peptide gene of the complete candidates concerning the constructed database (EFI-EST: 35% identity).

This methodology provides a valuable tool for the identification and prioritization of novel specialized metabolites, particularly in RiPPs, by enabling analysis at the single-gene level. The approach is expected to have significant implications for drug discovery research, as it facilitates the identification of previously unknown metabolites with potential therapeutic applications.

### Antimicrobial Peptide Prioritization

Recognizing the growing need for developing new antibiotics [49,50] and *Streptomyces*’ significant potential to produce them, particularly within the RiPP class, which demonstrates a remarkable ability to generate novel antimicrobial compounds [27,51], effective against ESKAPE pathogens [52-54], it becomes imperative to assess the potential antimicrobial activity in the identified candidates of potentially novel peptides obtained so far.

Five different machine-learning platforms were used to predict antimicrobial activity. Two of them showed the best area under the curve for both the precision-recall and the Receiver Operating Characteristic (ROC) curves. The default cutoff value of these platforms needed to be re-evaluated for RiPPs, since all platforms were trained with different experimental data and predictions. They were employed to identify the optimal similarity cutoff point, ensuring the best precision. Data collected from the RiPP manually constructed database was used to identify the best platform and cutoff value. Experimental information on their antimicrobial activity is available (Supplementary Table 1).

The generated ROC curve, plotting the true positive rate vs. the false positive rate, identifies the best model for discriminating between two classes, with a higher area under these curves being closer to 1. The two tools closest to 1 were MACREL, with a value of 0.68, and AmPEP, with a value of 0.64. Subsequently, the area under the precision-recall curve was analyzed and compared, with MACREL (0.89) and AmPEP (0.85) being the platforms closest to a value of 1. By plotting the precision and recall behavior over a range of cutoff values (0 to 1), MACREL exhibited almost constant data retrieval, allowing the selection of a cutoff value focused on achieving the highest precision without significantly affecting recall. Thus, a cutoff value 0.37 was selected to predict antimicrobial activity using the MACREL platform. It is important to note that the high “precision” values obtained in this analysis are likely overestimated due to a significant disparity in data between classes (AMP: 182 and Non-AMP: 52), directly impacting the analyses applied in this methodology.

Peptides were evaluated using their amino acid sequences as input for the MACREL platform, resulting in the prediction of 67 peptides with potential antimicrobial activity. Regarding their hemolytic potential, 24 were classified as hemolytic and 43 as non-hemolytic. The class distribution within these 67 peptides was highly diverse (see Fig. 4), with the predominant class being Class 1 lanthipeptides, totaling 21. This aligns with literature findings, where the substantial potential of lanthipeptides to produce bioactive compounds has been reported [55]. However, the notable contribution of Class V lanthipeptides is of great interest, given their recent discovery [56]. This emphasizes the strong potential of this class to produce new compounds with innovative biosynthetic machinery, confirming the viability of the methodology presented here for discovering a plethora of potentially novel compounds. Furthermore, it incorporates a strategy for identifying potentially novel small coding genes responsible for encoding a diverse array of compounds in the case of RiPPs. This is a critical limitation in current strategies, given the difficulty in accurately detecting and annotating small open reading frames [57].

**Fig.4.**
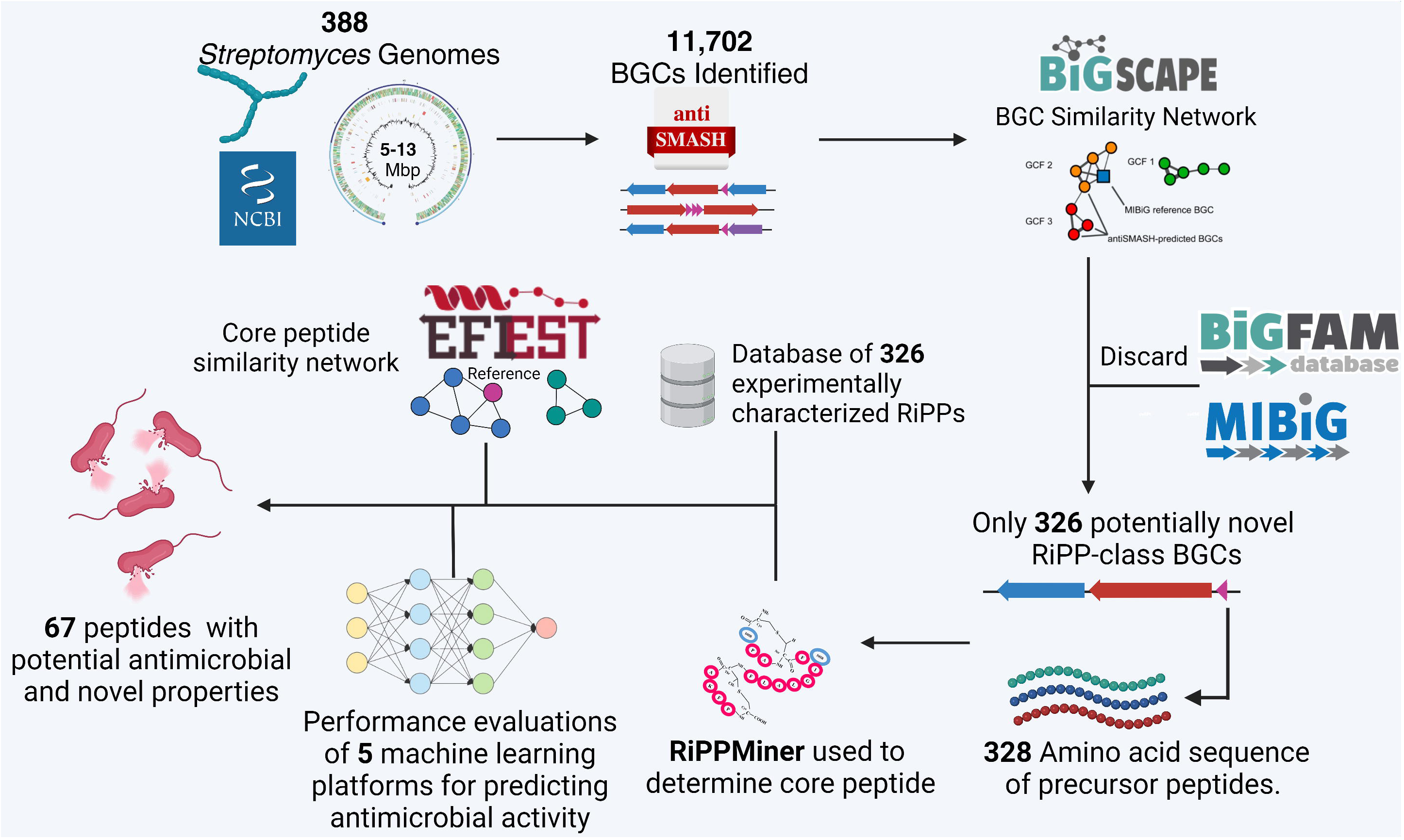
Distribution of novel candidate peptide classes. From the novel and complete BGCs, the possible antimicrobial activity was predicted using the MACREL AI with a cut-off value >0.37.

Our study has revealed significant genetic material in Streptomyces that remains unexplored for producing bioactive, specialized metabolites. To address this gap, we have developed a comprehensive methodology that prioritizes the discovery of novel natural products at three different levels: CGF, BGC, and the level of a single gene on a metagenomic scale. This methodology is achieved by implementing SSN tools such as BiG SCAPE/EFI-EST and comparing them with reference databases (BiG FAM/CGF, MiBIG/BGC, and a constructed database of experimentally characterized peptides). The primary focus of this methodology is to discover compounds with antimicrobial activity.

We have substantiated the success of this comprehensive methodology by discovering 67 peptides from Streptomyces with potential antimicrobial and novel properties. Therefore, using this methodology, considering the vast availability of genomic information can lead to relevant discoveries regarding the potential application of new compounds in clinical and various industrial settings.

In conclusion, our study highlights the importance of exploring the vast genetic material in Streptomyces to discover novel natural products. The methodology developed, and the discoveries made can pave the way for developing new antimicrobial agents with potential clinical and industrial applications.

## 8 Author statements

### 8.1 Author contributions

CDC, NSC, and SARA developed the idea and wrote and edited the manuscript. SARA and MAAH performed the bioinformatics analyses. MAAH wrote and edited the manuscript. We acknowledge using Grammarly AI to improve the manuscript’s English writing [58].

