## Supplementary Figure 1 for "Genome mining of *Streptomyces* for the discovery of low-resistance antibiotics"

**Suplementary Data 6.** results of the analysis of cuts in each AI used. The link to the script used is attached.
Scrip: https://github.com/Bert-IQ/ripps-classification

### Receiver Operating Characteristic (ROC) and AUC for each model

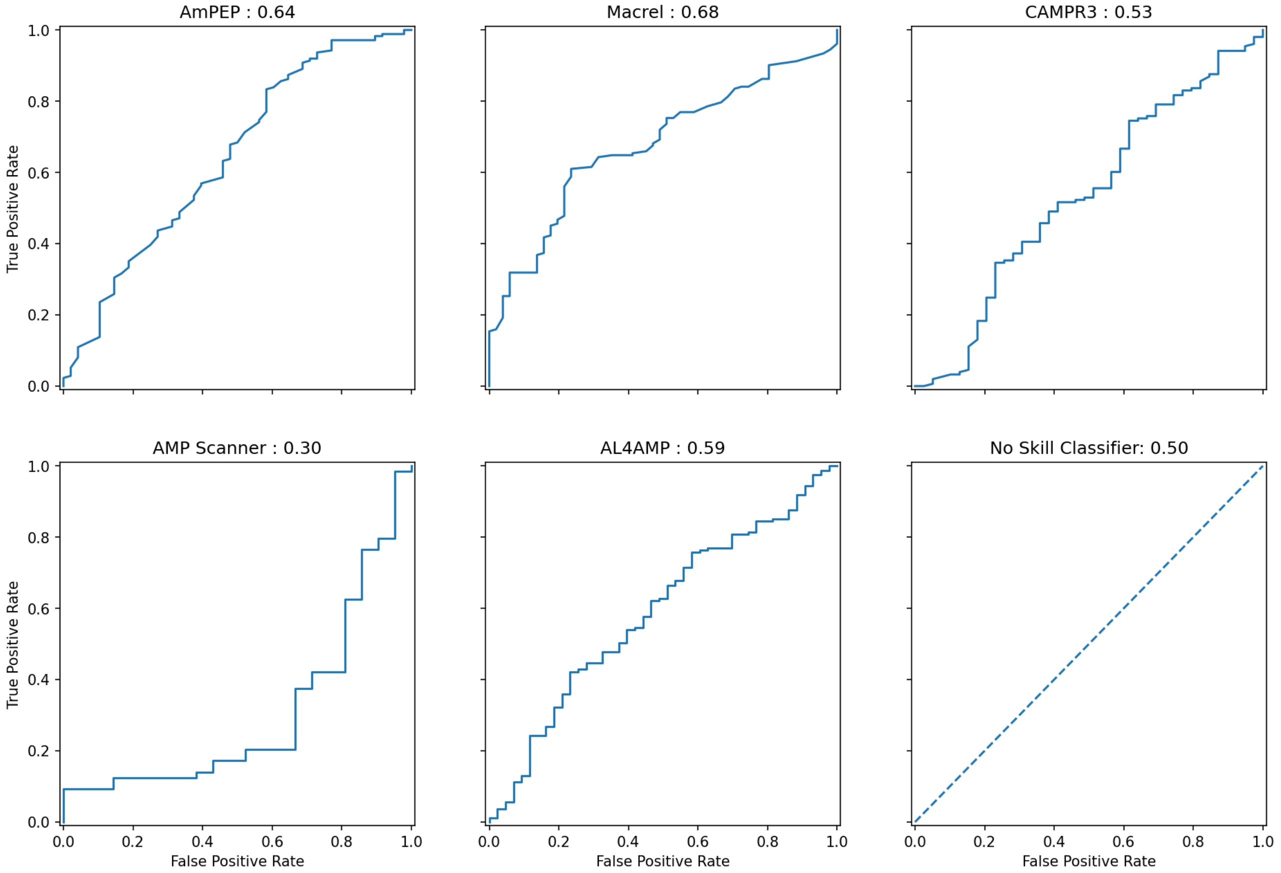


### PRC graphs and their AUC for each model.

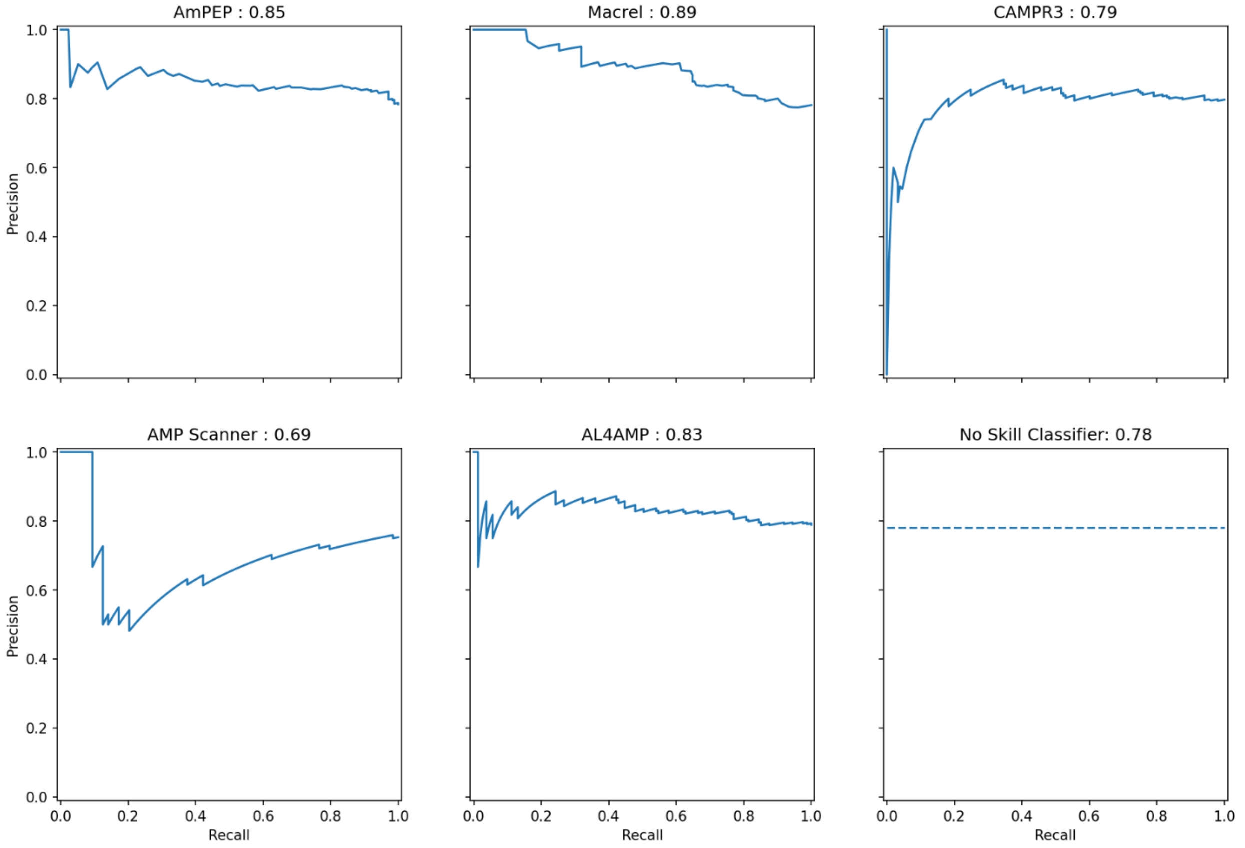

**More detailed analysis of the performance of AmPEP and Mackerel**

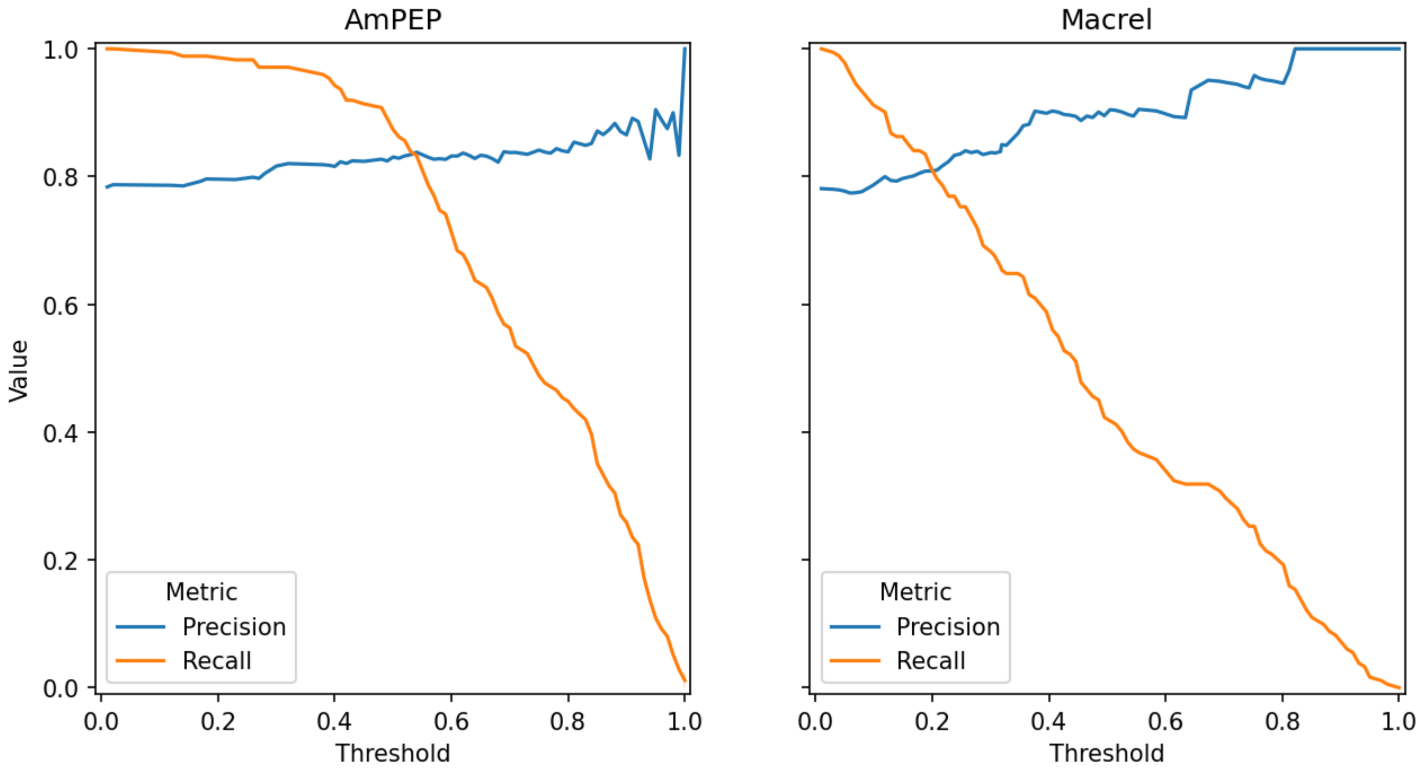
